# Viruses contribute to microbial diversification in the rumen ecosystem and are associated with certain animal production traits

**DOI:** 10.1101/2023.11.03.565476

**Authors:** Ming Yan, Zhongtang Yu

**Author notes:** Correspondence note: Zhongtang Yu, Department of Animal Sciences, The Ohio State University, Columbus, OH 43210, USA.

## Abstract

**Background:** The rumen microbiome enables ruminants to digest otherwise indigestible feedstuffs, thereby facilitating the production of high-quality protein, albeit with suboptimal efficiency and producing methane. Despite extensive research delineating associations between the rumen microbiome and ruminant production traits, the functional roles of the pervasive and diverse rumen virome remain to be determined.

**Results:** Leveraging a recent comprehensive rumen virome database, this study analyzes virus-microbe linkages, at both species and strain levels, across 551 rumen metagenomes, elucidating patterns of microbial and viral diversity, co-occurrence, and virus-microbe interactions. Additionally, this study assesses the potential role of rumen viruses in microbial diversification by analyzing prophages found in rumen metagenome-assembled genomes. Employing CRISPR-Cas spacer-based matching and virus-microbe co-occurrence network analysis, this study suggests that rumen viruses may regulate rumen microbes at both strain and community levels via both antagonistic and mutualistic interactions. Moreover, this study establishes that the rumen virome demonstrates responsiveness to dietary shifts and associations with key animal production traits, including feed efficiency, lactation performance, weight gain, and methane emissions.

**Conclusions:** These findings furnish a substantive framework for subsequent investigations to decode the functional roles of the rumen virome in shaping the rumen microbiome and influencing overall animal production performance.

## Introduction

The rumen microbiome plays an essential role in providing nutrients to ruminants by digesting fibrous feedstuffs that would otherwise remain indigestible and converting low-quality dietary nitrogen into high-quality microbial protein. Nonetheless, the microbial processes involved are inefficient from a ruminant nutritional perspective and contribute to the emissions of methane (CH_4_) and waste nitrogen as urea and ammonia. Extensive research has sought to elucidate the interactions within the rumen microbiome, focusing on its association with diet, host genetics, and host phenotype (as reviewed in Huws, Creevey (1)). Remarkably, except for a very few, all these studies have emphasized the roles of bacteria, archaea, protozoa, and fungi, leaving rumen viruses largely overlooked. Consequently, this has led to a paucity of knowledge concerning the ecological and nutritional roles, as well as significance, of rumen viruses (1), despite being the most numerically abundant entities in the rumen ecosystem (2) and acting as potential apex hierarchy regulators of the rumen microbiome and nutrient recycling.

Microbial viruses significantly impact microbiomes across diverse ecosystems. In ocean settings, viruses lyse approximately 20% of the microbes daily (3), profoundly influencing biogeochemical cycles through the enhancement of carbon and nitrogen recycling via “viral shunt” (4), a process that is modulated by virome diversity in a spatiotemporal manner (5, 6). In contrast, the human gut virome remains relatively stable (7) but displays considerable variation across an individual’s lifespan (8) and is linked to chronic diseases (9). Rumen viruses, both abundant(2) and diverse (10), infect diverse rumen microbes including the core rumen microbiome. By lysing rumen microbes at different trophic levels, rumen viruses likely regulate the populations and activities of their hosts and thus the recycling of nutrients and microbial protein (11), which serves as the primary metabolizable protein that ruminants utilize. Thus, unraveling the complexities of virus-microbe interactions within the rumen ecosystem is crucial for deciphering the implication of the rumen virome in animal production performance metrics, encompassing aspects such as feed efficiency, lactation performance, and CH_4_ emissions.

Viruses affect the diversity, population dynamics, and metabolic activities of various microbes through several hypothetical modes of virus-microbe interactions, such as “kill the winner”, “piggyback the winner”, and the “arms-races” dynamic (12). Specifically, by selectively lysing dominant microbial strains, viruses contribute to the maintenance of microbial diversity. They also facilitate host adaptation and diversification by facilitating horizontal gene transfer (13). Furthermore, viruses drive microbial diversification through adaptive co-evolution (14). Prophages, whether cryptic or non-cryptic, serve as accessory gene reservoirs that may carry genes enhancing host survival (15). Moreover, viruses can modulate host metabolism directly by expressing auxiliary metabolic genes (AMGs), thereby influencing critical ecological processes in both the environment and host-associated ecosystems, including the human gut and the rumen (10, 16).

Previous studies have documented variations in rumen virome in response to dietary shifts (17) and proposed their potential effects on nutritionally important rumen bacteria (2, 18, 19). However, the rumen virome remains poorly understood in terms of diversity, interactions with their hosts, and its roles in regulating rumen functions and animal production performance. To bridge this knowledge gap, we recently developed a comprehensive rumen virome database (RVD) by employing the latest bioinformatics pipelines for metagenomic virome analysis across nearly 1,000 rumen metagenomes (10). We revealed a vast diversity of viruses infecting various taxa of rumen bacteria, archaea, and protozoa within the rumen, along with a diverse repertoire of AMGs, including those encoding nutritionally essential enzymes such as cellulases. The revelation of these rumen viruses along with the linkages with their hosts implies a substantial influence on the rumen ecosystem. Building on these findings, we posited that the rumen virome interacts intimately with the rumen microbes across multiple paradigms and connects to important animal production traits including methane emissions. To test this hypothesis, we systematically characterized and analyzed the prophages at the strain level to decipher viral host specificity at both inter- and intra-species levels. Furthermore, we built and compared microbe-virus and microbe-only networks to ascertain the roles of rumen viruses in shaping rumen microbiome structure. Finally, we analyzed 311 rumen metagenomes reported in nine independent studies, uncovering associations between the rumen virome, microbiome, and critical animal production traits, including feed efficiency, lactation performance, and CH_4_ emissions. Collectively, the results demonstrate that the rumen virome plays pivotal roles in regulating rumen microbial assembly, diversification, and functions, and it is intricately connected with diet and several important animal production traits.

## Method

### Developing and benchmarking custom kraken2 classifiers tailored for the rumen microbiome

We developed three custom Kraken2 classifiers based on the GTDB taxonomy and utilizing three databases: the representative genomes of GTDB R207 (65,703 genomes), GTDB R207 plus 3,588 high-quality rumen MAGs (>90 complete, <5% contamination), and GTDB R207 plus 7,176 high-quality dereplicated MAGs assembled from rumen metagenomes in the present study (see supplementary information for details). We benchmarked these new Kraken2 classifiers against the standard Kraken2 classifier using the rumen metagenomic data reported by Xue, Wu (20), which were not used to assemble rumen MAGs or other analyses in the present study. The newly developed Kranken2 classifier that incorporated GTDB R207 and the 7,176 rumen MAGs, referred to as the Rumen Kraken2 Classifier, was used in further analysis.

### Species-level profiling and identification of “core” species of the rumen microbiome

We performed species-level profiling of the 975 rumen metagenomes described in the previous study (10) using the Rumen Kraken2 Classifier. The number of sequence reads assigned to individual species was computed using Bracken (21), with the outputs then being compiled and imported to R 4.0.2 (22). Only the species each represented by >0.001% of the total assigned reads were considered present in a sample. The prevalence of each species and genus was calculated, and the species and genera with a 100% prevalence were regarded as core/ubiquitous. The relative abundance of each species was calculated as the proportion of the reads assigned to that species relative to the sum of all taxonomically assigned reads. To investigate the influence of sample size on the identification of core species/genera, we employed a custom Python script. Briefly, starting with a randomly selected 100 samples, we incrementally added 5 random samples each iteration and re-calculated the counts of core species/genera, until all the 975 samples were included. This process was repeated 100 times, and the resulting counts of core species/genus across the iterations were plotted and visualized in R.

### Virome profiling and ecological analysis for alpha- and beta-diversity, differential abundance, and virus-to-host ratio

Of the original studies reporting the 975 metagenomes, nine reported comprehensive metadata, including details of experimental design, dietary treatments, and animal production metrics (Supplementary Table 1). We profiled the viromes within the metagenomes derived from these nine studies by mapping the quality-filtered reads to the RVD using CoverM (option: --min-read-percent-identity 0.95, --min-read-aligned-percent 0.75, --min-covered-fraction 0.7; https://github.com/wwood/CoverM) and the trimmed mean method. The number of reads mapped to the RVD per Gb of metagenomic reads was used as a proxy for viral richness. The Kruskal-Wallis Test in R was used to assess the statistical difference in viral richness among treatments or animal groups. We also calculated the corresponding microbial richness based on the microbes that were classified in each metagenome using the Rumen Kraken2 Classifier. Spearman correlation coefficients computed in R were used to identify correlations between viral and microbial richness.

We conducted beta-diversity analysis of the rumen viromes using PCoA, based on Bray-Curtis dissimilarity, through the vegan package (23) in R. To test for difference among treatments or animal groups, we performed PERMANOVA using the adonis2 function of the same package, with 999 permutations, testing for difference among treatments or animal groups. When comparing the rumen viromes among feed efficiencies in beef cattle, breed was considered a confounding factor, and PERMANOVA was performed with restricted permutations (“strata = breed”).

Differential abundance analysis of both the microbial and viral profiles across treatments or animal groups was conducted using LinDA (24). To exclude potentially spurious minor species, only those with a relative abundance exceeding 0.01% in at least 60% of the samples were included. Additionally, vOTUs found in fewer than 50% of the metagenomes were excluded. The resulting *p*-values were adjusted for multiple testing with the Benjamini– Hochberg procedure, and significance was declared at an adjusted *p*-value (*q*) < 0.1.

To assess whether variations in diet or animal production performance are associated with changes in prophage lifecycle, we computed the virus-to-host ratio (VHR), which is defined as the ratio of the prophage genomes coverage rate (determined by CoverM) to the number of reads assigned to their predicted host species (based on the Bracken result). We then compared the VHR across treatments or animal groups in nine studies, focusing on the virus-host linkages identified in at least six rumen metagenomes per treatment or animal group. We used the Kruskal-Wallis test to assess significance with the *p-*values adjusted for multiple testing using the Benjamini–Hochberg procedure in R.

### Identification, taxonomic classification, and host prediction of prophages identified in the rumen MAGs

Using VirSorter2 V2.2.3 (25), We identified the viral sequences from the 7,176 rumen MAGs used to develop the Rumen Kraken2 Classifier, the 1,726 RUG2 MAGs, and the Hungate1000 collection, as described in the previous study (10). The quality of these viral sequences was verified using CheckV 0.8.1 (26), and only those meeting the VirSorter2 category 1 and 2 criteria were retained. These viral sequences underwent further confirmation validation using VIBRANT (27) V1.2.1 (-virome), and only those confirmed again to be viral were retained as bona fide viral sequences. The confirmed viral sequences were further annotated using DRAM-v V1.2.4 (28). Two categories of viral sequences were identified as prophages: those flagged as integrated prophages (extracted from the host contigs) by CheckV, and those present in contigs that contain any of the prophage-related genes including integrase (VOG00021), excisionase (VOG00006, VOG05065), cro repressor (VOG00002), and cl repressor (VOG00692), as identified based on the VOGDB database (https://vogdb.org/). We then clustered the identified prophage sequences into vOTU at 95% average nucleotide identity (ANI) across at least 85% of the shortest contigs using scripts from CheckV (26) (https://bitbucket.org/berkeleylab/checkv/src/master/scripts/). The prophage vOTUs were taxonomically classified using PhaGCN2 (29), and their host linkages were inferred from the MAGs they were identified. A phylogenetic tree that includes the identified host species of the “core” genera was generated using GTDB-tk (option: -classify_wf) (30) and visualized using Itol (31). Active prophages, which display a significantly higher coverage than the flanking host genome regions (based on the reads mapping results), were identified with PropagAtE V1.1.0 (32). We then mapped the reads of the 975 metagenomes to the prophages that have an identified host. A prophage was assumed non-cryptic if it was predicted to be active by PropagAtE in any metagenomes.

### Identification of ARGs carried by prophages

We identified ARGs present in the prophage sequences using the stringent criteria established by Enault, Briet (33). Specifically, we predicted the ORFs of the prophage sequences using Prodigal (34) and then aligned them to the CARD 3.2.6 database. Genes conferring resistance via specific mutations were excluded, as recommended previously (35). Prophage contigs with a greater than 80% identity with a database sequence and 40% coverage were retained for manual curation, as described previously (10).

### Micro-diversity analysis of prophage-carrying strains

First, we identified the MAGs representing individual strains from the 7,176 rumen MAGs used to develop the Rumen Kraken2 Classifier and the 1,726 RUG2 MAGs using drep (--S_algorithm fastANI -- greedy_secondary_clustering -ms 10000 -pa 0.9 -sa 0.98 -nc 0.30 -cm larger). These representative MAGs were combined to form a ‘MAG mapping database’. To minimize read mis-mapping, we prioritized the RUG2 MAGs using the “--extra_weight_table” flag. Second, we profiled each RUG2 sample at the strain level by mapping its reads to the MAGs mapping database using InStrain (36). Third, we calculated the pN/pS ratio of individual genes within each strain detected in each metagenome. A strain was deemed present if at least five reads were mapped to at least 50% of its MAGs, as recommended previously (36). A strain was considered to carry prophage(s) when its scaffolds reach breadth >99%. We computed the pN/pS ratio for each prophage gene with criteria of greater than 99% breadth and 10X coverage. The prophage genomic structure was visualized with the gggenes package in R.

### Host prediction with CRISPR spacer matching at the strain level

We predicted the CRISPR-Cas arrays across the high-quality RUG2 MAGs (37) using MinCED (38). The identified spacer sequences, at least 30 bp, were then matched to the vOTUs identified from the RUG2 samples (extracted from the RVD) using BLASTn with a threshold of 100% sequence identity. The presence of MAGs and vOTUs in each RUG2 sample was examined using InStrain. We identified genome-level virus-host linkages requiring co-occurrence of both a MAG and a vOTU that have matching spacer sequences in the same RUG2 samples. The number of microbial strains without the spacer match but co-existing with strains of the same species with a spacer match was also determined. Only the linkages of sample ERR3275126 were visualized as a network using Cytoscape (39).

### Microbe-only and virus-microbe network analysis

Based on the microbial and viral profiles of the RUG2 samples, we constructed microbial-only and virus-microbe networks. To eliminate minor potentially spurious vOTUs and their hosts, we included only major microbial species with a relative abundance exceeding 0.01% in at least 70% of the samples and vOTUs with a trimmed mean (based on CoverM) value exceeding 1 in at least 50% of the samples. Both networks were constructed using SpiecEasi (40) with the sparse graphical lasso (glasso) setting, as described by Tipton, Müller (41). The networks were visualized in R with the package igraph (42). We computed the network modularity and assortativity with the “fastgreedy.community()” and “assortativity()” functions of igraph, respectively. We also analyzed the data at a 70% prevalence threshold. The degree centrality of microbial nodes was compared between the microbe-only and virus-microbe networks with one-tailed paired t-test in R.

## Results

### A custom rumen Kranken2 classifier tailored to the rumen microbiome enhances the classification and identification of rumen microbes

A custom Kraken2 classifier that incorporates the NCBI RefSeq complete genomes, the Hungate 1000 collection (43), and rumen metagenome-assembled genomes (MAGs) substantially improved the classification rate of rumen metagenomic sequences (44). However, it failed to classify most of the MAGs-represented rumen microbes to the species level. This limitation arises from its reliance on the NCBI taxonomy, which is inadequate to capture the burgeoning numbers of rumen MAGs and thus constraints polyphyletic groupings (45). To refine species-level identification of virus-microbe interactions, we developed three custom Kraken2 classifiers based on the GTDB taxonomy and utilizing three databases: the representative genomes of GTDB R207 (65,703 genomes), GTDB R207 plus 3,588 high-quality rumen MAGs (>90 complete, <5% contamination), and GTDB R207 plus 7,176 high-quality rumen MAGs (refer to Methods for details). Compared with the standard Kraken2 classifier, the newly developed Kraken2 classifier using GTDB R207 enhanced the species-level classification rate by approximately 55%. The Kraken2 classifier that incorporates GTDB R207 and the additional 7,176 rumen MAGs - henceforth termed the Rumen Kraken2 Classifier – elevated species-level classification by 58% (Supplementary Fig. 1a). With a species-level classification rate exceeding 75%, the Rumen Kraken2 Classifier enhances species-resolved microbiome profiling, thereby facilitating accurate analysis of virus-microbe linkages and interactions in the rumen ecosystem.

Numerous studies have identified prevalent rumen microbes at the genus level using 16S rRNA gene sequencing (43, 46, 47). To explore virus-microbe interactions and assess the effects of rumen viruses on dominant rumen microbial species, we reanalyzed the 975 rumen metagenomes analyzed in a previous study on rumen viromes (10). We discovered a set of ubiquitous species (100% prevalence), most of which belong to the genus *Prevotella* (Supplementary Fig. 2a). Notably, the combined relative abundance of the *Prevotella* species, reaching a relative abundance of 80% in some rumen metagenomes, was up to an order of magnitude higher than that of the next most prevalent genus, *Cryptobacteroides* (Supplementary Fig. 2b). Plotting the numbers of core species and genera against increasing sample size revealed a plateau at the species level but not at the genus level, indicating that inclusion of additional samples is unlikely to reduce the number of ubiquitous species identified (Supplementary Fig. 1b and 1c).

### Prophages are prevalent in the rumen ecosystem and may confer survival advantages to their hosts

The importance of lysogeny and the “piggyback the winner” model has been increasingly recognized in ecosystems densely populated by various microbes (48). To assess prophage prevalence in the rumen ecosystem, we comprehensively analyzed 8,902 rumen microbial genomes and MAGs and found 5,185 prophages that represent 4,225 vOTUs. Approximately 50% of these genomes and MAGs carry at least one prophage, with one MAG even carrying as many as eight prophages (Fig. 1a). The high prophage prevalence among the rumen microbial genomes/MAGs is comparable with that reported in bacteria in general (49). All the classifiable prophage vOTUs were classified under the order *Caudovirales*, with the majority of prevalent prophage vOTUs classified to families *Siphoviridae, Myoviridae*, and *Podoviridae*, while the less prevalent prophage vOTUs classifies to other families, including *Autographiviridae, Ackermannviridae*, and *Herelleviridae* (Fig. 1b). Of 5,185 identified prophages, 514 were predicted to be active (non-cryptic) in at least one sample. All 36 ubiquitous bacterial species examined for prophages presence were found to contain prophage sequences, and the majority carry both cryptic and non-cryptic prophages (Supplementary Fig. 3). The propensity for a host genome to carry non-cryptic prophages appeared to vary among bacteria, with the genomes from the phylum *Bacteroidota* more likely carrying non-cryptic prophages than those from *Firmicutes_A*.

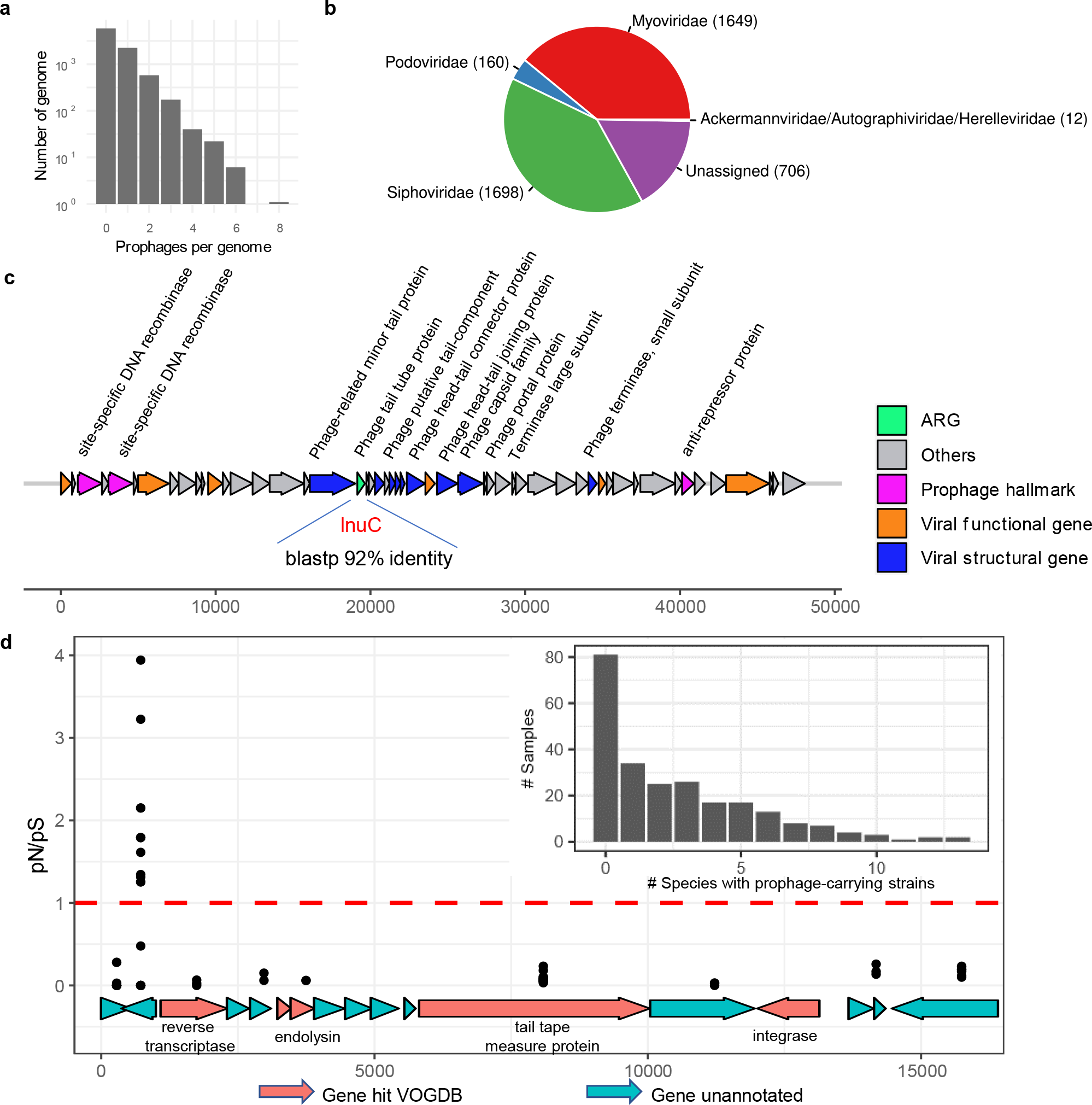

In a recent study, we identified antimicrobial resistance genes (ARGs) in some of the viral MAGs (10). The current study specifically targets ARGs carried by prophages. Our analysis revealed the presence of ARGs in multiple prophage genomes (Supplementary Table 2), including one prophage genome from a MAG of *Agathobacter sp900546625* (Fig. 1c). This particular prophage carries an ARG sharing 92% amino acids identity with *lnu*C, an ARG that confers resistance to lincomycin through nucleotidylation in *Streptococcus agalactiae* UCN36 (50). Since this ARG is demarcated by viral hallmark genes on both ends, it is unlikely part of the host genome. While most of the identified prophages were potentially cryptic, they may still confer adaptive advantages to their host, such as resistance to antibiotics, by providing accessory genes (15, 51, 52). The diversity and prevalence of these accessory genes, especially ARGs and genes involved in nutrient acquisition, warrants further investigation.

We further assessed the co-existence of multiple strains (individual MAGs) within the 240 samples/metagenomes used to derive the RUG2 catalog of rumen MAGs (44) (1,726 in total, referred to as ‘RUG2 MAGs’ hereafter). We found that most of the samples (hereafter referred to as ‘RUG2 samples’) had multiple species, each containing multiple prophage-carrying strains (Fig. 1d). Given that the MAG database only retains a limited subset of species for strain identification, the actual strain-level diversity is likely higher. Nevertheless, we found multiple strains of *Prevotella sp900317685*, including one strain carrying a cryptic prophage co-existing with other strains in 46 of the RUG2 samples. Examining the ratio of non-synonymous to synonymous substitutions (pN/pS, a measurement of gene micro-diversity), of the genes of this cryptic prophage, we found one unannotated gene with a pN/pS ratio exceeding one in most of these 46 samples (Fig. 1d), which indicates that this gene is undergoing positive diversifying selection. While the function of this gene is unknown, its presence may hint at survival advantages conferred by this prophage gene. Besides, temperate phages can also promote horizontal gene transfer (HGT) and microbial diversification not just through specialized and generalized transduction, the latter poof which is rare, but also through lateral transduction and conjugative transfer, both of which are common (14, 53). These processes can obscure the demarcation between host chromosomes and mobile genetic elements (53, 54, 55). Collectively, prophage-mediated HGT and the introduction of new genes during lysogenic conversion contribute to a beneficial relationship at the population level.

### Rumen viruses regulate microbiome at both species and strain levels

The intricate interplay between microbial defense mechanisms and viral countermeasures contributes to their co-evolution and shapes microbiome structure, especially at the strain level (56, 57). To explore these co-evolutionary dynamics, we examined the virus-host interactions across 1,422 high-quality MAGs and tens of thousands of vOTUs that we identified from the RUG2 samples. Employing CRISPR–Cas spacer matches, we assessed the co-existence and infection patterns between these MAGs and vOTUs at the strain level. We identified viruses with both inter- and intra-species host specificity, as exemplified by the virus-host linkages in one sample (Fig. 2a). Notably, many microbial genomes and MAGs contained CRISPR–Cas spacers that match multiple vOTUs. Because our analysis focused on high-quality MAGs, which are typically derived from highly abundant bacteria, the strain-level virus-host linkages we identified are likely skewed towards those abundant in the rumen ecosystem, such as strains of *Prevotella* species (e.g., *Prevotella sp900314935* and *Prevotella sp900314995*). Some MAGs had no match with the protospacer sequences of vOTUs predicted to infect the corresponding host species, indicating immunity or absence of previous infection. Since CRISPR spacers document past infections, concurrent detection of a matching CRISPR spacer and a protospacer in co-existing virus and microbe indicates a long-term coevolutionary relationship (58), which promotes both viral and microbial diversity. Moreover, we found many protospacers with a single mismatch with their corresponding CRISPR–Cas spacer. This could indicate point mutations in the viral genomes to evade the CRISPR-Cas system. The prevalence of CRISPR-Cas system among the rumen MAGs and genomes, together with previously identified restriction-modification systems (e.g., methyltransferase) in many rumen virus genomes (10), suggest that the “arms-races” model also plays a vital role in the rumen ecosystem. In analyzing the RUG2 samples, we found that about 80% of the vOTUs would infect just one single host strain, represented by one genome or MAG, and thus a single host species (Fig. 2b), while other vOTUs showed both inter- and intra-species host specificity. The broad host range of rumen viruses, as documented in previous studies (10, 59), may be attributable to, among others, mutations and rearrangements of receptor-binding proteins (60) and “sensitivity acquisition”, a process wherein bacteria initially resistant to phage infection become susceptible through receptor exchange with susceptible co-inhabitants (61). The strain-level microbial diversity in the rumen may be associated with host production traits. For example, in the initial investigation, methane emission was found not correlated to microbial abundance (62). However, subsequent research revealed that such a correlation existed at the strain level (44). Therefore, by regulating microbial community at both strain and species levels, rumen phages could also have an intricate relationship with animal production traits. Furthermore, the dynamic equilibrium between microbial defense and viral counter defense may result in oscillation in clonal abundance as a result of the genetic sweeps (63). Overall, the complex nested infections (phages infecting multiple strains/species and microbes infected by multiple phages) underscore the intricate virus-microbe interactions, which is further illustrated in the next section, and signify an important role of viruses in promoting trophic cascades as posited in a previous study (64).

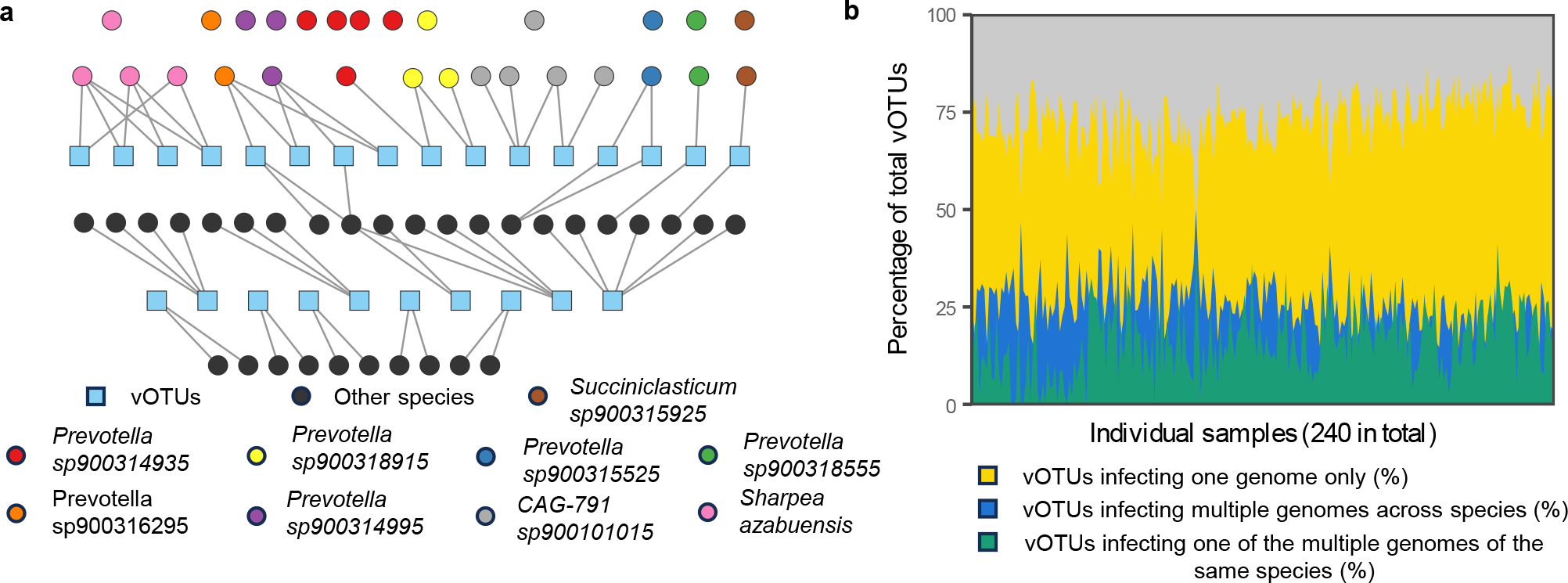

### Rumen viruses facilitate microbial interactions, as shown by virus-microbe networks

To investigate microbial interactions, we constructed a microbial co-occurrence network using the RUG2 samples. This network contains 671 microbial nodes, 119 of which are not linked to the main network (Fig. 3a). With an average degree centrality of 3.13 (± 3.24) and a modularity index of 0.71, the network displays a robust community structure. The network comprises three large, highly interconnected modules or discrete clusters of nodes. Each module has over 45 nodes, suggesting niche differentiation. We noted a moderate assortment among nodes based on their phyla (assortment coefficient c_a_ = 0.43). The largest module comprises 109 nodes, including primarily species within the genera of *Bacteroidota*, followed by species within genera of *Firmicutes, Firmicutes_A, Firmicutes_C, Fibrobacter, Actinobacteriota*, and archaea. The second largest module encompasses 93 nodes, mostly core species of *Prevotella* and *UBA4334* (a genomic genus in the family *Bacteroidaceae* in GTDB). This module also contains several genera of *Firmicutes_C* and *Proteobacteria* and archaea. The smallest module has 47 modes and features a diverse array of species from multiple phyla, including *Firmicutes, Firmicutes_A, Firmicutes_C, Actinobacteriota*, and *Bacteroidota* and archaea. All three modules contain unclassified species, indicating that some rumen microbes are not represented by the current GTDB database. Although the modules have the same set of phyla, they each have distinct genera, implying niche differentiation at finer taxonomic scales.

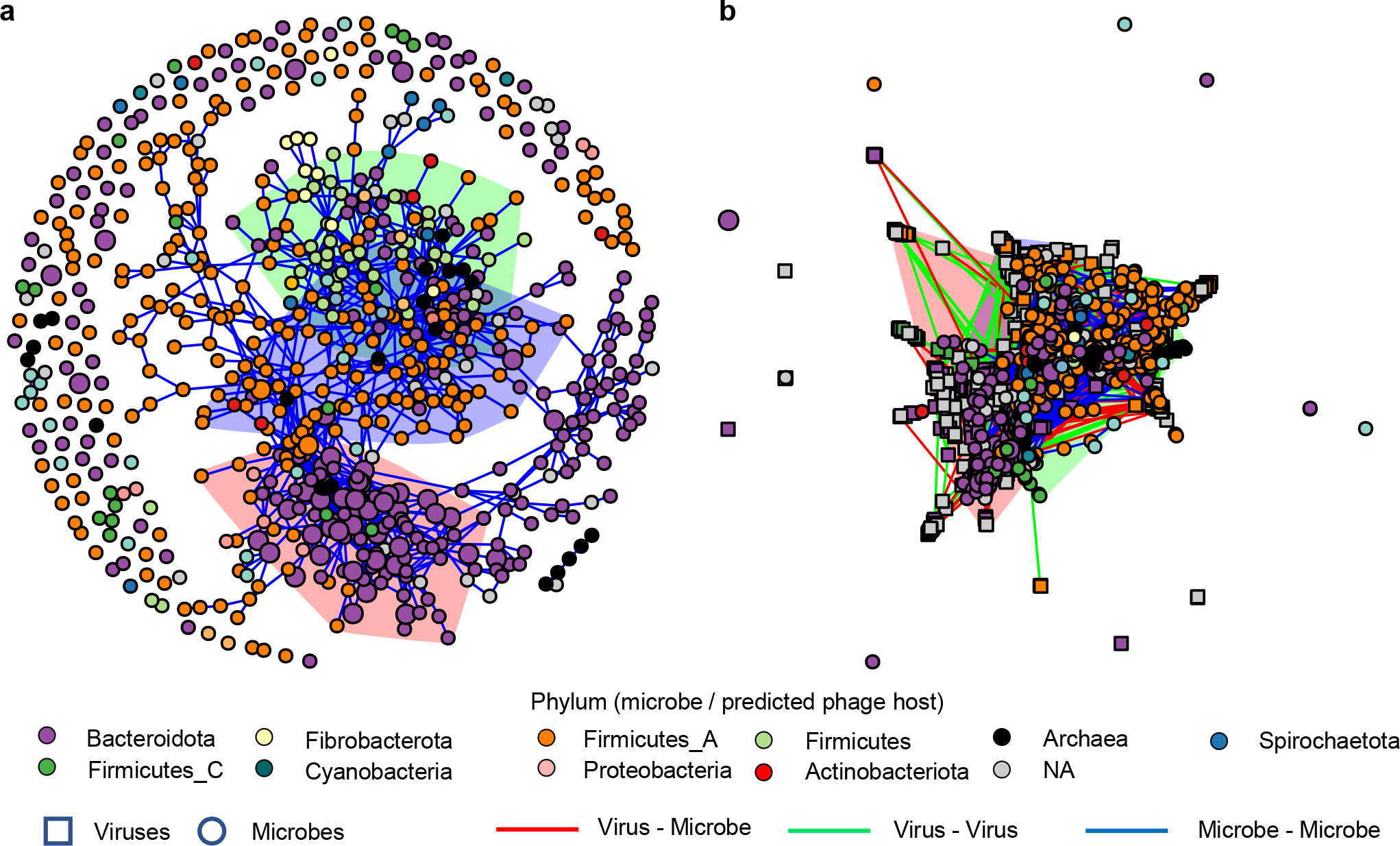

We also constructed a virus-microbe cooccurrence network to examine virus-microbe interactions (Fig. 3b). This network includes 570 viral nodes and the 671 microbial nodes of the microbe-only network. In this network, 22 microbial nodes do not connect to other microbial nodes. When considering only the microbial nodes, the average degree centrality is 5.23 (± 3.94), significantly higher than that of the microbe-only network (paired t-test, *p* < 0.001). With a modularity index of 0.60, relatively lower compared to that of the microbe-only network, the virus-microbe network still reveals a relatively robust community structure. Unlike in the microbe-only network, the three largest microbial modules in the virus-microbe network have a similar taxonomy composition (Supplementary Fig. 4a), and each contains multiple microbe-virus and virus-virus edges (Supplementary Fig. 4b). Moreover, the three modules are less separated (assortment coefficient c_a_ = 0.34) compared to microbe-only network. The microbe-virus edges can signify co-existence strategies, either as prophages within host microbes or as lytic viruses alongside virus-resistant microbes. In the latter scenario, it may be because virus-resistant microbes benefit from increased nutrient availability due to decreased competition and nutrients released from the microbes lysed by the lytic viruses, as shown previously (65). Although viruses may act antagonistically at the cell level, the augmented connectivity and reduced assortativity of the microbial nodes in the virus-microbe network, relative to the microbe-only network, suggest that viruses facilitate microbial interactions and allow diverse microbes to occupy the same niches. This inference is further supported by the modular and nested virus-microbe infection network as shown previously (Fig. 2a). Overall, these intricate virus-microbe interactions extend beyond the predator-prey relationship and indicate that rumen viruses and microbes could be mutualistic at the microbiome level, corroborating the previous finding in the human gut ecosystem (53). Interactions between phages can arise from superinfection immunity induced by prophages or co-infection of the same species. Repeating the analysis with an increased prevalence threshold from 50% to 70%, we noted increased connectivity and decreased assortativity (Supplementary Fig. 5). This indicates that the initial prevalence threshold did not bias the results.

Several microbial and viral nodes exhibit both a high degree centrality (>15) and a betweenness centrality (>15,000), as shown in Supplementary Fig. 6. These nodes include *Prevotella sp902778255, Prevotella sp900319305, GCA-900199385 sp017512985, UBA1711 sp001543385, RUG572 sp902802945*, and *Schwartzia succinivorans*. These species could be viewed as “keystone” species, crucial for maintaining community structure. Some of the nodes contain ubiquitous species but with a lower average degree centrality and betweenness centrality, 10 and 3,600, respectively. Modularity analysis suggests that while most of these ubiquitous species occupy distinct and essential niches, they may not be keystone species. Notably, two keystone viral species were predicted to infect *Ruminococcus_E sp900314795* and *CAG-791 sp900101015*. Given the intricate interplay between microbes and viruses, future rumen microbiome research should concurrently analyze both entities to understand the inconsistent and transient effects of microbial interventions reported in a previous study (66). Moreover, stochastic events affect the early colonization of the rumen and have a lasting influence on the rumen microbiome (67), but viruses were not taken into account. Future studies on the rumen ecosystem development should analyze rumen viruses.

### Dietary composition, animal production performance, and CH_4_ emissions are linked to the macro- and micro-diversity profiles of the rumen virome

The interrelationship between the rumen microbiome, diet, animal production performance, and CH_4_ emissions represents a key focus of rumen microbiome research. However, studies examining the connections of the rumen virome with the above factors or production traits remain scarce. Only one study has shown that dietary energy levels can affect both the rumen virome and microbiome (17). In the current study, we analyzed the rumen virome profiles of 311 rumen metagenomes across 9 studies, each with a detailed experimental design, to investigate the association between the rumen virome, diet, and animal production traits (Supplementary Table 1). To mitigate variability arising from differences in diet and animal genetics across the studies, we analyzed the data on a study-by-study basis. Overall, dietary composition affected virome richness, though the effect varied (Fig. 4a). For instance, beef cattle fed high-concentrate diets had a lower virome richness compared to those fed medium-concentrate diets. In dairy cattle, high lipid and starch diets corresponded to increased virome richness, while grazing led to a lower richness compared to total mixed ration (TMR, primarily consisting of corn silage and corn grain). Non-fiber carbohydrate (NFC) levels did not affect rumen virome richness in goats, but the levels of dietary protein and neutral detergent fiber (NDF, representing cellulose, hemicellulose, and lignin of plant fiber) appeared influential. Diets likely affect the rumen virome indirectly by affecting their hosts.

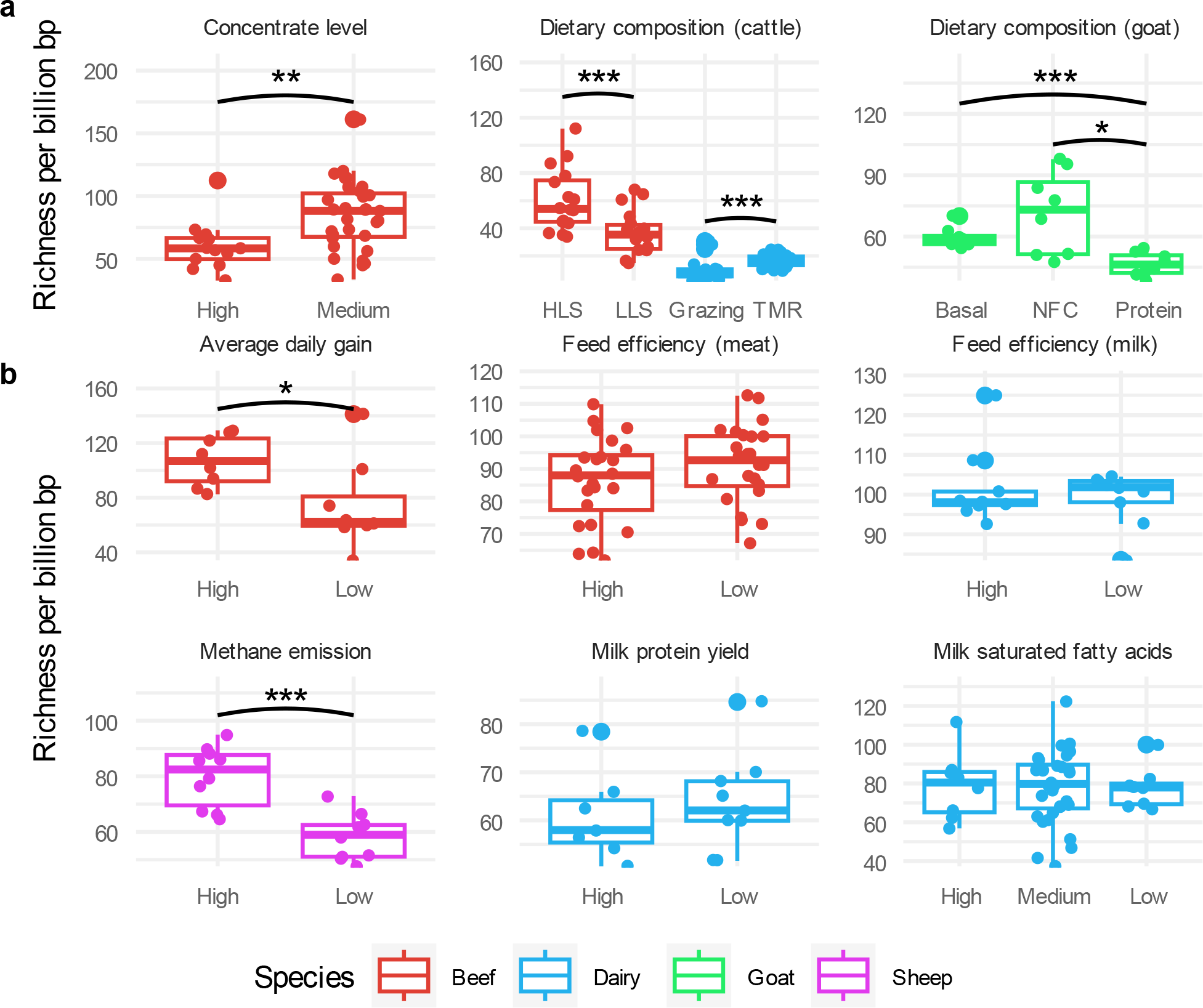

Average daily gain in beef cattle and CH_4_ emissions from sheep correlated positively with rumen viral richness; but feed efficiency in both beef cattle and dairy cows, as well as milk protein yield and saturated fatty acid yield, showed no association with virome richness (Fig. 4b). Animal production performance is affected by diet and a wide range of host factors such as age, metabolism, physiology, and health (68, 69, 70). The lack of significant association between the rumen virome and these animal production performance metrics may be attributable to those host factors. In examining the correlation between microbial richness and viral richness in the same rumen metagenomes, we found inconsistent results (Supplementary Fig. 7). Specifically, a significant correlation between microbiome and virome richness was observed only in some of the metagenomes, with no consistent directionality, suggesting that other factors likely affect their interactions and population dynamics. Given the highly individualized nature of the rumen virome (10, 53), interactions between rumen viruses and microbes, especially those at low abundance, may be affected by stochasticity constrained by the deterministic effects of diet and the host’s genetics.

We further analyzed the viral beta-diversity among animal groups using principal coordinates analysis (PCoA) based on Bray-Curtis dissimilarity. Particularly, rumen virome composition differed between diets (Fig. 5a), feed efficiencies in beef cattle (breeds as a confounding factor), CH_4_ emissions from sheep, and milk protein yields. Conversely, no differences were observed among average daily gains in beef cattle, feed efficiencies, or milk saturated fatty acid yields in lactating dairy cows (Fig. 5b). Although some studies have reported correlations between rumen microbiome composition and the above animal production traits (62, 71, 72, 73), other studies have not (74). The divergence in the association between animal production performance and the rumen virome, relative to the rumen microbiome, may be attributable to the more individualized rumen virome profiles than the microbiome profiles.

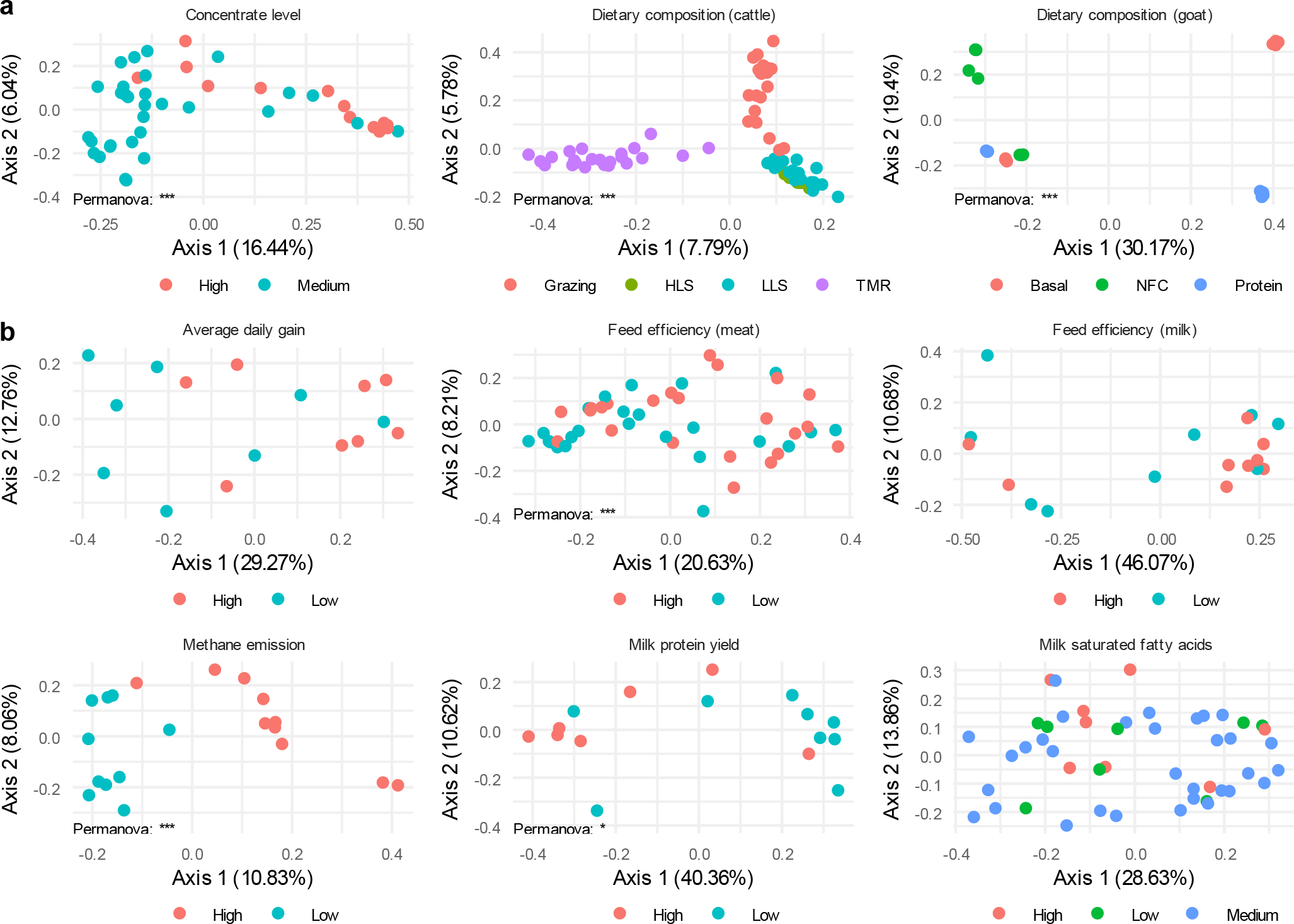

### Rumen viruses affect microbial species depending on dietary conditions and animal production performance

In addition to modifying microbiome structure, rumen viruses may directly modulate rumen fermentation by affecting the abundance of key microbial species. Using differential abundance analysis, we identified several dozens of vOTUs with different abundance (*q* < 0.1) across varying dietary compositions (Fig. 6a), and animal production metrics (Fig. 6b). Because a considerable proportion of the vOTUs could not be classified at any taxonomy rank above vOTUs, differential abundance was analyzed only at this granularity. The hosts of some differentially abundant vOTUs also displayed varied abundance (*q* < 0.1; indicated by red arrows in Fig. 6). Notably, vOTU FH88564_121008||full, predicted to infect *Prevotella brevis*, was more prevalent in the medium concentrate group, whereas *Prevotella brevis* itself was more prevalent in the high concentrate group. Conversely, vOTU, ERR3275101_45023||full and its predicted host, *Succiniclasticum sp900315925*, exhibited the same trend: more prevalent in the low CH_4_ emission group than in the high CH_4_ emission group. Since these two vOTUs were predicted to be prophages, their divergent trends may signify disparate life cycles. The hosts of some vOTUs could not be predicted, probably because their abundance was below the 0.01% threshold. None of the AMG-encoding vOTUs was differentially abundant, possibly due to their limited representation in the dataset used in the analysis.

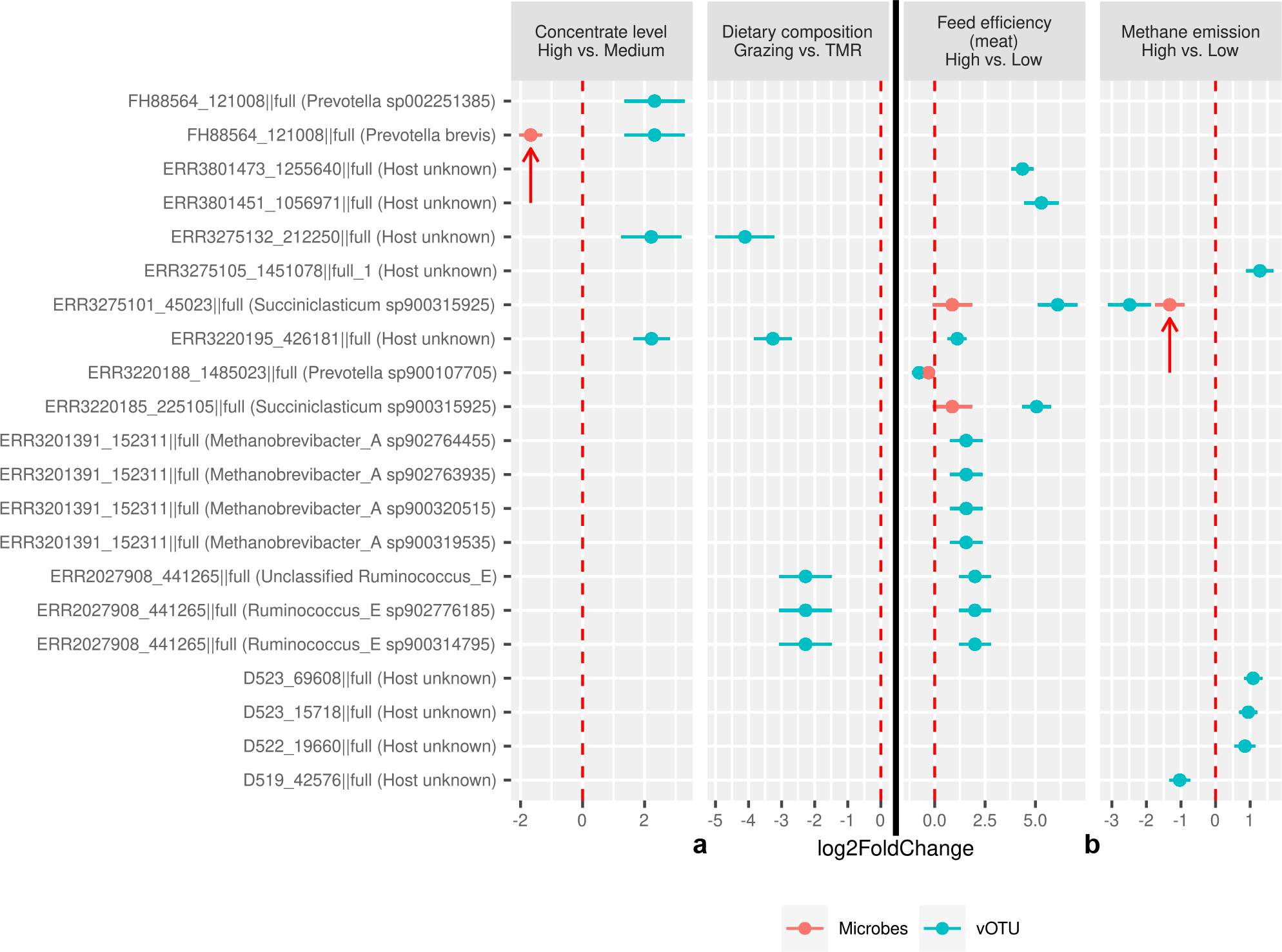

We evaluated potential associations between diet or animal production performance and lifecycle alterations of prophages by comparing the VHR among the predicted virus-host linkages across the 311 rumen metagenomes used for diversity analysis. We found a higher VHR (*q <* 0.01) for the prophages predicted to infect *Prevotella sp002251295* and *Prevotella sp900107705* in the animals fed a concentrate-based diet (NFC) compared with those fed a forage-based diet (Supplementary Fig. 8a). This disparity is likely attributable to the increased feed fermentation and production of short-chain fatty acids (SCFAs), which are known to induce prophages (75). Indeed, certain food and food extracts have been shown to induce prophages in the human microbiome (76). Dietary fructose and SCFAs also potentiated prophage induction in *Lactobacillus reuteri*, a gut microbe (75). Additionally, subacute rumen acidosis, generally induced by rapid SCFA production in animals consuming high-concentrate diets, has been shown to substantially increase rumen viral abundance (10). Thus, although shifts in the rumen virome largely mirror alterations in microbiome structure, changes in the rumen environment may modulate viral lifecycle dynamics, which can in turn affect the rumen microbiome structure. Intriguingly, the VHR between prophage vOTU FH88564_121008||full and its host, *Prevotella brevis*, remained unaffected by the concentrate levels, despite their differential abundance at the two concentrate levels. This can likely be attributed to the concurrent presence of multiple strains of the host species, and they do not carry the same prophage, as shown in the previous section. Moreover, sheep with varying CH_4_ emissions exhibited significantly different VHR (Supplementary Fig. 8b), which may be ascribed to alterations in viral lifecycle dynamics induced by shifts in microbial metabolisms (62).

The turnover of rumen microbes caused by viral lysis can have a far-reaching effect on certain rumen functions, especially fermentation and microbial protein synthesis. Although marine phages are estimated to lyse approximately 20% of the marine bacteria daily (3), the lysis rate of rumen microbes attributable to viruses remains undetermined. Given the high abundance of both viruses and microbes in the rumen, viral lysis therein is likely substantial. Two key questions thus arise: What is the virus-mediated turnover rate of both total and specific rumen microbes, particularly those pertinent to animal production performance and CH_4_ emissions? And to what extent do lysogenic and lytic cycles predominate in the rumen ecosystem? Early studies used transmission electron microscopy to count phages (2) or total phage DNA concentration as a proxy of phage population size in the rumen (18). However, a high phage count does not necessarily correlate with an elevated host mortality rate. For example, less than 1% of the cyanobacterial cells were infected even in the presence of high concentrations of free phage particles (77). It is worth noting that the above two methods likely underestimate phage abundance because they primarily account for free lytic virions. In contrast, VHR calculated from metagenomic sequences can quantify not only free virions but also temperate phages and intracellular lytic phages (78). Furthermore, the single-cell polony method can help identify lineage-resolved viral infections across thousands of cells of various microbes simultaneously (77). Leveraging these methodological advancements, future studies should aim to quantify the virus-host ratio, host mortality rate, and their associations with net microbial protein synthesis in the rumen ecosystem. Such data will provide invaluable insights into the role of viral lysis in intra-ruminal nitrogen cycling across different feeding regimes.

## Conclusions

In conclusion, this study delves into the largely uncharted territory of the rumen virome’s role within the rumen microbiome. Through the examination of comprehensive rumen metagenomes, as well as the genetic material of both microbes and viruses, it reveals intricate virus-microbe relationships, providing insights into the diversity, co-occurrence, and interactions between these components. Furthermore, the study suggests that rumen viruses may exert regulatory influence on rumen microbes at both strain and community levels through both antagonistic and mutualistic interactions. Notably, the rumen virome displays adaptability in response to dietary changes and exhibits associations with crucial animal production traits, such as feed efficiency, lactation performance, weight gain, and methane emissions. These findings establish a robust foundation for future research endeavors aimed at deciphering the functional roles of the rumen virome in shaping the rumen microbiome and its profound impact on overall animal production performance.

## References

1. Huws SA, Creevey CJ, Oyama LB, Mizrahi I, Denman SE, Popova M, et al. Addressing global ruminant agricultural challenges through understanding the rumen microbiome: past, present, and future. Frontiers in microbiology. 2018;9:2161.

2. Ritchie A, Robinson I, Allison M. Rumen bacteriophage: survey of morphological types. Microscopie electronique. 1970;3:333–4.

3. Suttle CA. Marine viruses—major players in the global ecosystem. Nature reviews microbiology. 2007;5(10):801–12.

4. Breitbart M, Bonnain C, Malki K, Sawaya NA. Phage puppet masters of the marine microbial realm. Nature microbiology. 2018;3(7):754–66.

5. Hevroni G, Flores-Uribe J, Béjà O, Philosof A. Seasonal and diel patterns of abundance and activity of viruses in the Red Sea. Proceedings of the National Academy of Sciences. 2020;117(47):29738–47.

6. Brum JR, Hurwitz BL, Schofield O, Ducklow HW, Sullivan MB. Seasonal time bombs: dominant temperate viruses affect Southern Ocean microbial dynamics. The ISME journal. 2016;10(2):437–49.

7. Shkoporov AN, Clooney AG, Sutton TD, Ryan FJ, Daly KM, Nolan JA, et al. The human gut virome is highly diverse, stable, and individual specific. Cell host & microbe. 2019;26(4):527–41. e5.

8. Guidi L, Chaffron S, Bittner L, Eveillard D, Larhlimi A, Roux S, et al. Plankton networks driving carbon export in the oligotrophic ocean. Nature. 2016;532(7600):465–70.

9. Tisza MJ, Buck CB. A catalog of tens of thousands of viruses from human metagenomes reveals hidden associations with chronic diseases. Proc Natl Acad Sci U S A. 2021;118(23).

10. Yan M, Pratama AA, Somasundaram S, Li Z, Jiang Y, Sullivan MB, et al. Interrogating the viral dark matter of the rumen ecosystem with a global virome database. Nature Communications. 2023;14(1):5254.

11. Gilbert RA, Townsend EM, Crew KS, Hitch TC, Friedersdorff JC, Creevey CJ, et al. Rumen virus populations: technological advances enhancing current understanding. Frontiers in microbiology. 2020;11:450.

12. Brown TL, Charity OJ, Adriaenssens EM. Ecological and functional roles of bacteriophages in contrasting environments: marine, terrestrial and human gut. Current Opinion in Microbiology. 2022;70:102229.

13. Dion MB, Oechslin F, Moineau S. Phage diversity, genomics and phylogeny. Nature Reviews Microbiology. 2020;18(3):125–38.

14. Mangalea MR, Duerkop BA. Fitness trade-offs resulting from bacteriophage resistance potentiate synergistic antibacterial strategies. Infection and immunity. 2020;88(7):10.1128/iai.00926-19.

15. Wang X, Kim Y, Ma Q, Hong SH, Pokusaeva K, Sturino JM, et al. Cryptic prophages help bacteria cope with adverse environments. Nature communications. 2010;1(1):147.

16. Kieft K, Breister AM, Huss P, Linz AM, Zanetakos E, Zhou Z, et al. Virus-associated organosulfur metabolism in human and environmental systems. Cell reports. 2021;36(5):109471.

17. Anderson CL, Sullivan MB, Fernando SC. Dietary energy drives the dynamic response of bovine rumen viral communities. Microbiome. 2017;5(1):155.

18. Klieve AV, Swain RA, Nolan J. Natural variability and diurnal fluctuation of bacteriophage populations in the rumen. 1993.

19. Friedersdorff JC, Kingston-Smith AH, Pachebat JA, Cookson AR, Rooke D, Creevey CJ. The isolation and genome sequencing of five novel bacteriophages from the rumen active against Butyrivibrio fibrisolvens. Frontiers in microbiology. 2020;11:1588.

20. Xue MY, Wu JJ, Xie YY, Zhu SL, Zhong YF, Liu JX, et al. Investigation of fiber utilization in the rumen of dairy cows based on metagenome-assembled genomes and single-cell RNA sequencing. Microbiome. 2022;10(1):11.

21. Lu J, Breitwieser FP, Thielen P, Salzberg SL. Bracken: estimating species abundance in metagenomics data. PeerJ Computer Science. 2017;3:e104.

22. R Core Team R. R: A language and environment for statistical computing. R foundation for statistical computing Vienna, Austria; 2018.

23. Oksanen J, Kindt R, Legendre P, O’Hara B, Stevens MHH, Oksanen MJ, et al. The vegan package. Community ecology package. 2007;10(631-637):719.

24. Zhou H, He K, Chen J, Zhang X. LinDA: linear models for differential abundance analysis of microbiome compositional data. Genome biology. 2022;23(1):1–23.

25. Guo J, Bolduc B, Zayed AA, Varsani A, Dominguez-Huerta G, Delmont TO, et al. VirSorter2: a multi-classifier, expert-guided approach to detect diverse DNA and RNA viruses. Microbiome. 2021;9(1):1–13.

26. Nayfach S, Camargo AP, Schulz F, Eloe-Fadrosh E, Roux S, Kyrpides NC. CheckV assesses the quality and completeness of metagenome-assembled viral genomes. Nature biotechnology. 2021;39(5):578–85.

27. Kieft K, Zhou Z, Anantharaman K. VIBRANT: automated recovery, annotation and curation of microbial viruses, and evaluation of viral community function from genomic sequences. Microbiome. 2020;8(1):1–23.

28. Shaffer M, Borton MA, McGivern BB, Zayed AA, La Rosa SL, Solden LM, et al. DRAM for distilling microbial metabolism to automate the curation of microbiome function. Nucleic acids research. 2020;48(16):8883–900.

29. Jiang J-Z, Yuan W-G, Shang J, Shi Y-H, Yang L-L, Liu M, et al. Virus classification for viral genomic fragments using PhaGCN2. Briefings in Bioinformatics. 2023;24(1):bbac505.

30. Chaumeil PA, Mussig AJ, Hugenholtz P, Parks DH. GTDB-Tk: a toolkit to classify genomes with the Genome Taxonomy Database. Bioinformatics. 2019.

31. Letunic I, Bork P. Interactive Tree Of Life (iTOL) v4: recent updates and new developments. Nucleic acids research. 2019;47(W1):W256–W9.

32. Kieft K, Anantharaman K. Deciphering active prophages from metagenomes. Msystems. 2022;7(2):e00084–22.

33. Enault F, Briet A, Bouteille L, Roux S, Sullivan MB, Petit MA. Phages rarely encode antibiotic resistance genes: a cautionary tale for virome analyses. Isme J. 2017;11(1):237–47.

34. Hyatt D, Chen GL, Locascio PF, Land ML, Larimer FW, Hauser LJ. Prodigal: prokaryotic gene recognition and translation initiation site identification. BMC Bioinformatics. 2010;11:119.

35. Alcock BP, Raphenya AR, Lau TT, Tsang KK, Bouchard M, Edalatmand A, et al. CARD 2020: antibiotic resistome surveillance with the comprehensive antibiotic resistance database. Nucleic acids research. 2020;48(D1):D517–D25.

36. Olm MR, Crits-Christoph A, Bouma-Gregson K, Firek BA, Morowitz MJ, Banfield JF. inStrain profiles population microdiversity from metagenomic data and sensitively detects shared microbial strains. Nature Biotechnology. 2021;39(6):727–36.

37. Stewart RD, Auffret MD, Warr A, Wiser AH, Press MO, Langford KW, et al. Assembly of 913 microbial genomes from metagenomic sequencing of the cow rumen. Nature communications. 2018;9(1):1–11.

38. Bland C, Ramsey TL, Sabree F, Lowe M, Brown K, Kyrpides NC, et al. CRISPR recognition tool (CRT): a tool for automatic detection of clustered regularly interspaced palindromic repeats. BMC Bioinformatics. 2007;8(1):1–8.

39. Shannon P, Markiel A, Ozier O, Baliga NS, Wang JT, Ramage D, et al. Cytoscape: a software environment for integrated models of biomolecular interaction networks. Genome research. 2003;13(11):2498–504.

40. Kurtz ZD, Müller CL, Miraldi ER, Littman DR, Blaser MJ, Bonneau RA. Sparse and compositionally robust inference of microbial ecological networks. PLoS computational biology. 2015;11(5):e1004226.

41. Tipton L, Müller CL, Kurtz ZD, Huang L, Kleerup E, Morris A, et al. Fungi stabilize connectivity in the lung and skin microbial ecosystems. Microbiome. 2018;6:1–14.

42. Csardi G, Nepusz T. The igraph software package for complex network research. InterJournal, complex systems. 2006;1695(5):1–9.

43. Seshadri R, Leahy SC, Attwood GT, Teh KH, Lambie SC, Cookson AL, et al. Cultivation and sequencing of rumen microbiome members from the Hungate1000 Collection. Nature biotechnology. 2018;36(4):359–67.

44. Stewart RD, Auffret MD, Warr A, Walker AW, Roehe R, Watson M. Compendium of 4,941 rumen metagenome-assembled genomes for rumen microbiome biology and enzyme discovery. Nat Biotechnol. 2019;37(8):953–61.

45. Parks DH, Chuvochina M, Waite DW, Rinke C, Skarshewski A, Chaumeil PA, et al. A standardized bacterial taxonomy based on genome phylogeny substantially revises the tree of life. Nat Biotechnol. 2018;36(10):996–1004.

46. Henderson G, Cox F, Ganesh S, Jonker A, Young W, Janssen PH. Rumen microbial community composition varies with diet and host, but a core microbiome is found across a wide geographical range. Scientific reports. 2015;5(1):14567.

47. Wallace RJ, Sasson G, Garnsworthy PC, Tapio I, Gregson E, Bani P, et al. A heritable subset of the core rumen microbiome dictates dairy cow productivity and emissions. Science advances. 2019;5(7):eaav8391.

48. Knowles B, Silveira C, Bailey B, Barott K, Cantu V, Cobián-Güemes A, et al. Lytic to temperate switching of viral communities. Nature. 2016;531(7595):466–70.

49. Touchon M, Bernheim A, Rocha EP. Genetic and life-history traits associated with the distribution of prophages in bacteria. The ISME journal. 2016;10(11):2744–54.

50. Achard A, Villers C, Pichereau V, Leclercq R. New lnu (C) gene conferring resistance to lincomycin by nucleotidylation in Streptococcus agalactiae UCN36. Antimicrobial agents and chemotherapy. 2005;49(7):2716–9.

51. Duerkop BA, Clements CV, Rollins D, Rodrigues JL, Hooper LV. A composite bacteriophage alters colonization by an intestinal commensal bacterium. Proceedings of the National Academy of Sciences. 2012;109(43):17621–6.

52. Bossi L, Fuentes JA, Mora G, Figueroa-Bossi N. Prophage contribution to bacterial population dynamics. Journal of bacteriology. 2003;185(21):6467–71.

53. Shkoporov AN, Turkington CJ, Hill C. Mutualistic interplay between bacteriophages and bacteria in the human gut. Nature Reviews Microbiology. 2022;20(12):737–49.

54. Huang J, Dai X, Wu Z, Hu X, Sun J, Tang Y, et al. Conjugative transfer of streptococcal prophages harboring antibiotic resistance and virulence genes. The ISME Journal. 2023:1–15.

55. Humphrey S, Fillol-Salom A, Quiles-Puchalt N, Ibarra-Chávez R, Haag AF, Chen J, et al. Bacterial chromosomal mobility via lateral transduction exceeds that of classical mobile genetic elements. Nature communications. 2021;12(1):6509.

56. Hampton HG, Watson BN, Fineran PC. The arms race between bacteria and their phage foes. Nature. 2020;577(7790):327–36.

57. Pilosof S, Alcala-Corona SA, Wang T, Kim T, Maslov S, Whitaker R, et al. The network structure and eco-evolutionary dynamics of CRISPR-induced immune diversification. Nature Ecology & Evolution. 2020;4(12):1650–60.

58. Stern A, Mick E, Tirosh I, Sagy O, Sorek R. CRISPR targeting reveals a reservoir of common phages associated with the human gut microbiome. Genome research. 2012;22(10):1985–94.

59. Bickhart DM, Watson M, Koren S, Panke-Buisse K, Cersosimo LM, Press MO, et al. Assignment of virus and antimicrobial resistance genes to microbial hosts in a complex microbial community by combined long-read assembly and proximity ligation. Genome biology. 2019;20(1):1–18.

60. de Jonge PA, Nobrega FL, Brouns SJ, Dutilh BE. Molecular and evolutionary determinants of bacteriophage host range. Trends in microbiology. 2019;27(1):51–63.

61. Tzipilevich E, Habusha M, Ben-Yehuda S. Acquisition of phage sensitivity by bacteria through exchange of phage receptors. Cell. 2017;168(1):186–99. e12.

62. Shi W, Moon CD, Leahy SC, Kang D, Froula J, Kittelmann S, et al. Methane yield phenotypes linked to differential gene expression in the sheep rumen microbiome. Genome research. 2014;24(9):1517–25.

63. Held NL, Herrera A, Cadillo-Quiroz H, Whitaker RJ. CRISPR associated diversity within a population of Sulfolobus islandicus. PLOS ONE. 2010;5(9):e12988.

64. Koskella B, Brockhurst MA. Bacteria–phage coevolution as a driver of ecological and evolutionary processes in microbial communities. FEMS Microbiology Reviews. 2014;38(5):916–31.

65. Daly RA, Roux S, Borton MA, Morgan DM, Johnston MD, Booker AE, et al. Viruses control dominant bacteria colonizing the terrestrial deep biosphere after hydraulic fracturing. Nature microbiology. 2019;4(2):352–61.

66. Ban Y, Guan LL. Implication and challenges of direct-fed microbial supplementation to improve ruminant production and health. J Anim Sci Biotechnol. 2021;12(1):109.

67. Furman O, Shenhav L, Sasson G, Kokou F, Honig H, Jacoby S, et al. Stochasticity constrained by deterministic effects of diet and age drive rumen microbiome assembly dynamics. Nature communications. 2020;11(1):1–13.

68. Li F, Li C, Chen Y, Liu J, Zhang C, Irving B, et al. Host genetics influence the rumen microbiota and heritable rumen microbial features associate with feed efficiency in cattle. Microbiome. 2019;7(1):1–17.

69. Xue M-Y, Sun H-Z, Wu X-H, Liu J-X. Multi-omics reveals that the rumen microbiome and its metabolome together with the host metabolome contribute to individualized dairy cow performance. Microbiome. 2020;8(1):1–19.

70. Xue M, Sun H, Wu X, Guan LL, Liu J. Assessment of rumen microbiota from a large dairy cattle cohort reveals the pan and core bacteriomes contributing to varied phenotypes. Applied and environmental microbiology. 2018;84(19):e00970–18.

71. Xue MY, Xie YY, Zhong Y, Ma XJ, Sun HZ, Liu JX. Integrated meta-omics reveals new ruminal microbial features associated with feed efficiency in dairy cattle. Microbiome. 2022;10(1):32.

72. Li F, Hitch TC, Chen Y, Creevey CJ, Guan LL. Comparative metagenomic and metatranscriptomic analyses reveal the breed effect on the rumen microbiome and its associations with feed efficiency in beef cattle. Microbiome. 2019;7(1):1–21.

73. Wallace RJ, Rooke JA, McKain N, Duthie C-A, Hyslop JJ, Ross DW, et al. The rumen microbial metagenome associated with high methane production in cattle. BMC Genomics. 2015;16(1):1–14.

74. Myer PR, Smith TP, Wells JE, Kuehn LA, Freetly HC. Rumen microbiome from steers differing in feed efficiency. PLOS ONE. 2015;10(6):e0129174.

75. Oh J-H, Alexander LM, Pan M, Schueler KL, Keller MP, Attie AD, et al. Dietary fructose and microbiota-derived short-chain fatty acids promote bacteriophage production in the gut symbiont Lactobacillus reuteri. Cell Host & Microbe. 2019;25(2):273–84. e6.

76. Boling L, Cuevas DA, Grasis JA, Kang HS, Knowles B, Levi K, et al. Dietary prophage inducers and antimicrobials: toward landscaping the human gut microbiome. Gut microbes. 2020;11(4):721–34.

77. Mruwat N, Carlson MC, Goldin S, Ribalet F, Kirzner S, Hulata Y, et al. A single-cell polony method reveals low levels of infected Prochlorococcus in oligotrophic waters despite high cyanophage abundances. The ISME Journal. 2021;15(1):41–54.

78. López-García P, Gutiérrez-Preciado A, Krupovic M, Ciobanu M, Deschamps P, Jardillier L, et al. Metagenome-derived virus-microbe ratios across ecosystems. The ISME Journal. 2023:1–12.

